# Mutations in genes *lpxL1, bamA* and *pmrB* impair the susceptibility of cystic fibrosis strains of *Pseudomonas aeruginosa* to murepavadin

**DOI:** 10.1101/2023.10.09.561565

**Authors:** Aya Ghassani, Pauline Triponney, Maxime Bour, Patrick Plesiat, Katy Jeannot, MucoMicrobes study Group

## Abstract

Murepavadin is a peptidomimetic exhibiting specific inhibitory activity against *Pseudomonas* species. In the present study, its *in vitro* activity was assessed on 230 cystic fibrosis (CF) strains of *P. aeruginosa* isolated from twelve French hospitals, in comparison with twelve other antipseudomonal antibiotics. Although murepavadin is still in pre-clinical stage of development, 9.1% (*n*=21) of the strains displayed a resistance superior to 4 mg/L, a level at least 128-fold higher than the modal MIC value of the whole collection (≤ 0.06 mg/L). Whole-genome sequencing of these 21 strains along with more susceptible isogenic counterparts coexisting in the same patients revealed diverse mutations in genes involved in the synthesis (*lpxL1* and *lpxL2*) or transport of lipopolysaccharides (*bamA, lptD*, and *msbA*), or encoding histidine kinases of two-component systems (*pmrB* and *cbrA*). Allelic replacement experiments with wild-type reference strain PAO1 confirmed that alteration of genes *lpxL1, bamA* and/or *pmrB* can increase murepavadin resistance from 8- to 32-fold. Furthermore, we found that specific amino-acid substitutions in histidine kinase PmrB (G188D, Q105P, and D45E) reduce the susceptibility of *P. aeruginosa* to murepavadin, colistin and tobramycin, three antibiotics used or intended to be used (murepavadin) in aerosols to treat colonized CF patients. Whether colistin or tobramycin may select mutants resistant to murepavadin or the opposite needs to be addressed by clinical studies.

## Introduction

*Pseudomonas aeruginosa* is a major cause of morbidity and mortality in cystic fibrosis (CF) (1). Because of its ability to survive in multiple environments, this Gram-negative pathogen frequently colonizes the airways of CF individuals generating *in situ* a chronic inflammation itself responsible for a decline of the respiratory function. In an attempt to control such a deleterious lung invasion, international guidelines recommend the administration of repeated cures of inhaled antibiotics to chronically infected patients (2, 3). Aerosols of tobramycin, colistin methane sulfonate, and in a lesser extent aztreonam are thus commonly used with this indication. More recently, murepavadin (POL7080, Spexis), a new peptidomimetic derived from the porcine cationic antimicrobial peptide protegrin-I secreted by neutrophils, has been recognized as potentially useful to treat CF and non-CF bronchiectasis patients by inhalation (upcoming Phase 1 clinical trial) (4, 5). The project of using the intravenous route was abandoned because of significant nephrotoxic effects (6). This 14 amino acid-long, β-hairpin-configured, cationic macrocyclic peptide that is stabilized by a D-proline-L-proline bond, is selectively active on *Pseudomonas* species (4). Unlike colistin which interacts with negatively charged residues born by the lipid A of lipopolysaccharides (LPS), murepavadin targets outer membrane proteins, mainly the β-barrel LPS transport protein D (LptD) (7, 8). This latter forms a complex with the outer membrane anchored protein LptE, to translocate newly synthetized LPS molecules from the periplasmic space to the bacterial surface. Interaction of murepavadin with or near the β-jellyroll periplasmic domain of LptD is believed to prevent the correct insertion of LPS into the outer membrane, leading to detrimental misfunctions (9). In preclinical studies, murepavadin showed an excellent *in vitro* activity (MIC_90_ from 0.12 to 0.25 mg/L) on non-CF clinical strains of *P. aeruginosa*, some of those being multidrug resistant (10, 11). Despite the potential application of the peptide in CF, its activity on CF strains was documented rather scarcely while revealing bacteria with MIC values greater than 4 mg/L (12). Because *P. aeruginosa* can adapt easily to most antipseudomonal antibiotics through mutations (13), some studies focused on the emergence of resistant mutants to murepavadin or its peptidomimetic analogue POL7001 *in vitro* (14-16). Thus, tandem duplication of a sequence LRDKGM in protein LptD was associated with a 64-fold increased resistance of reference strain PAO1 (4); whereas Romano et *al*. found that complementation of strain PA14 with *pmrB* alleles encoding altered peptides (G185S, G188D, and L172del) resulted in 2- to 16-fold higher murepavadin MICs (16). Finally, alteration of several genes (*cbrA, acrB2, lpxL1, lpxL2, lpxlT, msbA*, and *bamA*) by missense or frameshift mutations was predicted to reduce murepavadin susceptibility of PAO1 or its hypermutator mutant PAO1Δ*mutS* in time-kill experiments (15). However, except for some *pmrB* mutants, the impact of these mutations on murepavadin activity was not confirmed further.

The present study was set up to improve our knowledge on the antipseudomonal activity of murepavadin. Its MIC values were compared to that of currently used antibiotics for 230 CF isolates collected from 105 patients in 12 French hospitals. To get an insight into the mechanisms contributing to a decreased activity of the peptide in this particular clinical context, we next compared the genomic sequences of isolates coexisting in a same patient but differing in their resistance levels, and introduced the most prevalent mutations found into wild-type reference *P. aeruginosa* strain PAO1. Thus, we show that some of these mutations generate a cross-resistance between murepavadin and common antibiotics in CF such as tobramycin and colistin. The risk of co-selection of multidrug resistant strains with murepavadin in CF needs to be considered, especially in a hypermutator genetic background.

## Results and discussion

### In vitro susceptibility of CF strains to murepavadin

Two hundred and thirty isolates of *P. aeruginosa* collected over a three-month period from 105 French CF patients were tested for their resistance levels to 13 antipseudomonal antibiotics including murepavadin (Table 1 and Table S1). According to the EUCAST 2023 breakpoints established for *P. aeruginosa*, 10.9% of these isolates were susceptible or susceptible at increased exposure to all the currently approved antibiotics (no breakpoints have been defined yet for murepavadin), 38.7% were non-susceptible to at least one agent in less than three antimicrobial categories, 37.0% fitted with the definition of MDR, 12.1% were XDR, and 1.3% PDR (17). Among these molecules, colistin (94.4%), ceftazidime plus avibactam (87.4%), ceftolozane plus tazobactam (80.0%), and meropenem (80.0%) were the most frequently active (Table 1). The MIC values of murepavadin ranged from ≤ 0.06 to ≥ 128 mg/L (Table 1). While the murepavadin MIC_50_ value determined on our collection (0.125 mg/L) was identical to that reported previously for non-CF strains, a notable proportion of CF strains appeared to be more resistant (MIC_90_ = 4 mg/L) than the isolates of these studies (0.12 and 0.25 mg/L, respectively) (10, 11). Corroborating this observation, the MIC_90_ of the antibiotic was found equal to 2 mg/L for CF strains collected in Northern Ireland, the Netherlands, Spain, and Australia (12). Because murepavadin is intended to be administrated to CF patients by aerosolization, its activity was compared to that of antibiotics already used under the form of aerosols, such as colistin, aztreonam-lysin, and tobramycin. MIC_50_/MIC_90_ values of these molecules were equal to 1/2 mg/L, 4/128 mg/L and 2/32 mg/L, respectively (Table 1). Murepavadin retained a good activity on most strains considered as clinically resistant to tobramycin (MIC > 2 mg/L), colistin (MIC > 4mg/L) and aztreonam (MIC > 16 mg/L), thereby suggesting that this new drug could be an alternative to these common treatments. On the other hand, a high resistance to the peptide (arbitrarily fixed > 4 mg/L) was noted in 16 (7%), 8 (3.5%) and 9 (3.9%) isolates resistant to the three antibiotics respectively. Finally, a few strains with such relatively high murepavadin MICs turned out to be susceptible to one or more of these older molecules, mostly tobramycin (*n* = 5) and colistin (*n* = 13) (Table S1).

**TABLE 1.**
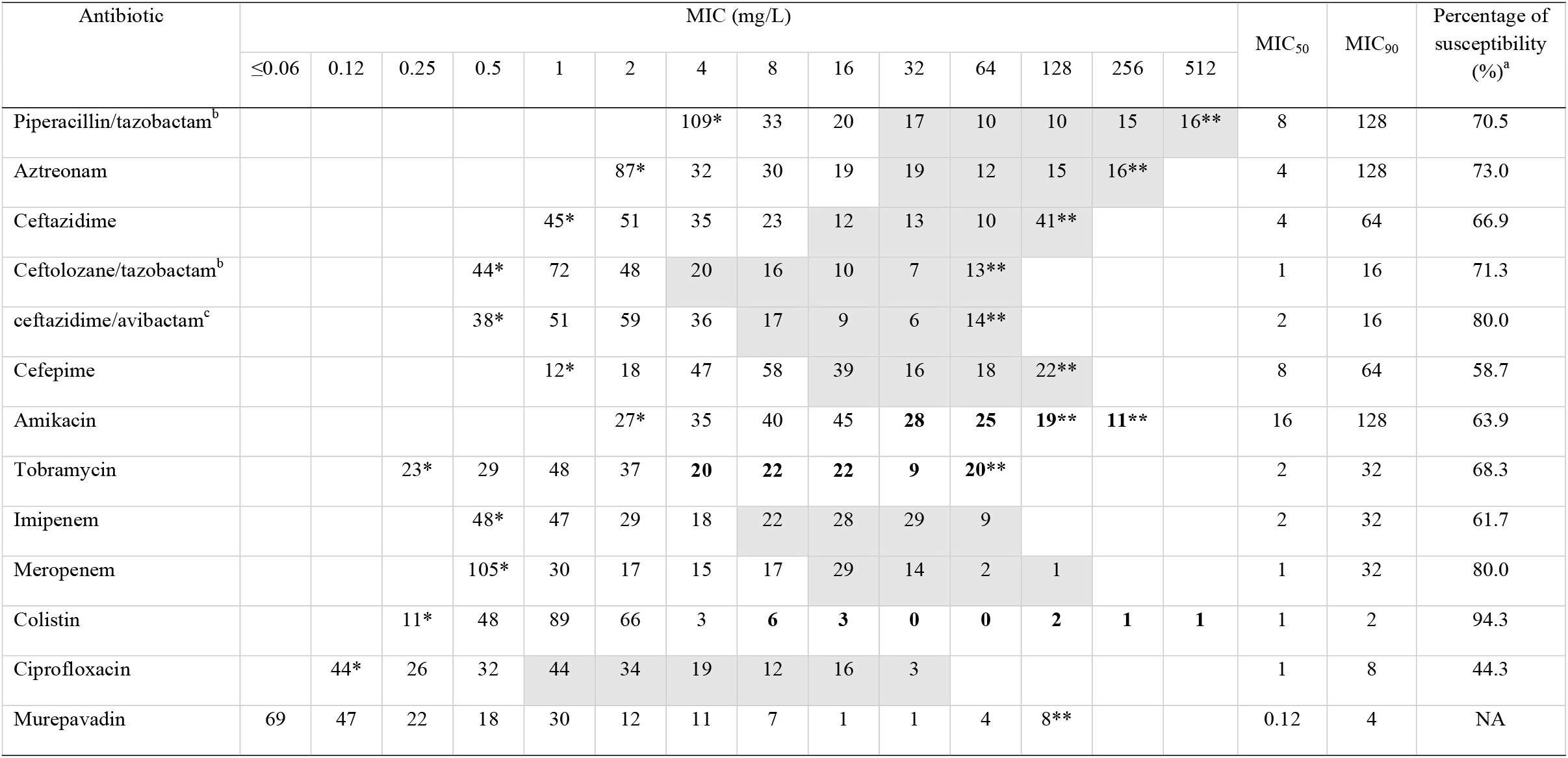
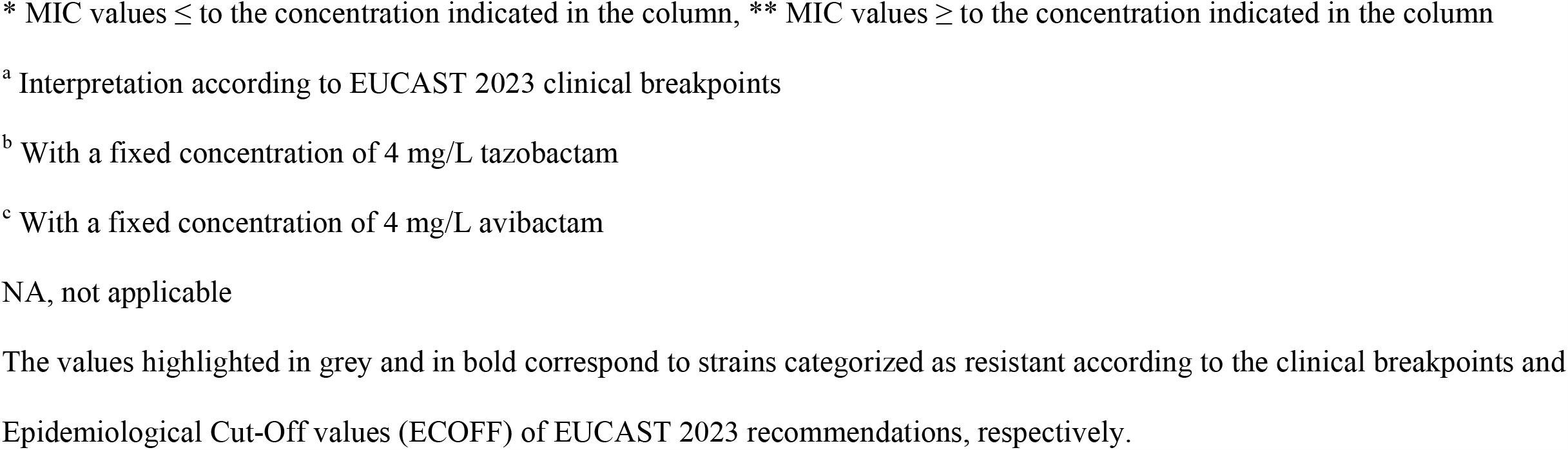
Antibiotic susceptibility levels of the 230 CF strains of *P. aeruginosa* selected for this study.

| Antibiotic | MIC (mg/L) |  |  |  |  |  |  |  |  |  |  |  |  |  | MIC<sub>50</sub> | MIC<sub>90</sub> | Percentage of susceptibility (%)<sup>a</sup> |
| --- | --- | --- | --- | --- | --- | --- | --- | --- | --- | --- | --- | --- | --- | --- | --- | --- | --- |
|  | ≤0.06 | 0.12 | 0.25 | 0.5 | 1 | 2 | 4 | 8 | 16 | 32 | 64 | 128 | 256 | 512 |  |  |  |
| Piperacillin/tazobactam<sup>b</sup> |  |  |  |  |  |  | 109* | 33 | 20 | 17 | 10 | 10 | 15 | 16** | 8 | 128 | 70.5 |
| Aztreonam |  |  |  |  |  | 87* | 32 | 30 | 19 | 19 | 12 | 15 | 16** |  | 4 | 128 | 73.0 |
| Ceftazidime |  |  |  |  | 45* | 51 | 35 | 23 | 12 | 13 | 10 | 41** |  |  | 4 | 64 | 66.9 |
| Ceftolozane/tazobactam<sup>b</sup> |  |  |  | 44* | 72 | 48 | 20 | 16 | 10 | 7 | 13** |  |  |  | 1 | 16 | 71.3 |
| ceftazidime/avibactam<sup>c</sup> |  |  |  | 38* | 51 | 59 | 36 | 17 | 9 | 6 | 14** |  |  |  | 2 | 16 | 80.0 |
| Cefepime |  |  |  |  | 12* | 18 | 47 | 58 | 39 | 16 | 18 | 22** |  |  | 8 | 64 | 58.7 |
| Amikacin |  |  |  |  |  | 27* | 35 | 40 | 45 | 28 | 25 | 19** | 11** |  | 16 | 128 | 63.9 |
| Tobramycin |  |  | 23* | 29 | 48 | 37 | 20 | 22 | 22 | 9 | 20** |  |  |  | 2 | 32 | 68.3 |
| Imipenem |  |  |  | 48* | 47 | 29 | 18 | 22 | 28 | 29 | 9 |  |  |  | 2 | 32 | 61.7 |
| Meropenem |  |  |  | 105* | 30 | 17 | 15 | 17 | 29 | 14 | 2 | 1 |  |  | 1 | 32 | 80.0 |
| Colistin |  |  | 11* | 48 | 89 | 66 | 3 | 6 | 3 | 0 | 0 | 2 | 1 | 1 | 1 | 2 | 94.3 |
| Ciprofloxacin |  | 44* | 26 | 32 | 44 | 34 | 19 | 12 | 16 | 3 |  |  |  |  | 1 | 8 | 44.3 |
| Murepavadin | 69 | 47 | 22 | 18 | 30 | 12 | 11 | 7 | 1 | 1 | 4 | 8** |  |  | 0.12 | 4 | NA |

\* MIC values ≤ to the concentration indicated in the column, \*\* MIC values ≥ to the concentration indicated in the column
<sup>a</sup> Interpretation according to EUCAST 2023 clinical breakpoints
<sup>b</sup> With a fixed concentration of 4 mg/L tazobactam
<sup>c</sup> With a fixed concentration of 4 mg/L avibactam
NA, not applicable
The values highlighted in grey and in bold correspond to strains categorized as resistant according to the clinical breakpoints and
Epidemiological Cut-Off values (ECOFF) of EUCAST 2023 recommendations, respectively.

Though none of the CF-patients from this work ever received murepavadin, 21 isolates (9.1%) from 15 individuals exhibited a resistance level greater than 4 mg/L, including eight isolates (3.5%) for which the MIC values were ≥ 128 mg/L (Table S1). To get an insight into the mechanisms involved in these phenotypes, we sequenced the genomes of these 21 bacteria along with those of more susceptible isolates (MIC ≤ 4 mg/L) coexisting in the same sputum samples (13 patients out of 15). Finally, intra-patient comparisons of these genome sequences were carried out in search of the most common SNPs (Table 2). The number of genomic alterations between concomitant clones varied from 2 (patient III-9) to 892 (patient XII-2) (Table S2). CF patients are often initially colonized by a single strain of *P. aeruginosa* which diversifies over the course of the disease to give rise to phenotypically distinct but genotypically related subpopulations well adapted to the lung environment (18). This evolution is usually boosted by the emergence of hypermutator clones deficient in one or several DNA proofreading systems (19, 20). Consistent with the relatively high divergence observed between some intra-patient clones, mutations in the DNA mismatch repair system (genes *mutS, mutL, uvrD*) and/or 8-oxodG system (genes *mutM, mutT, mutY*) were noticed in 23 out of the 37 sequenced strains (62.2%) (Table S2). On the other hand, a minimal divergence of 22 SNPs was found associated with a large murepavadin MIC difference of 1,024-fold (from 0.125 to ≥ 128 mg/L) between two clones colonizing patient XI-4.

**TABLE 2.**
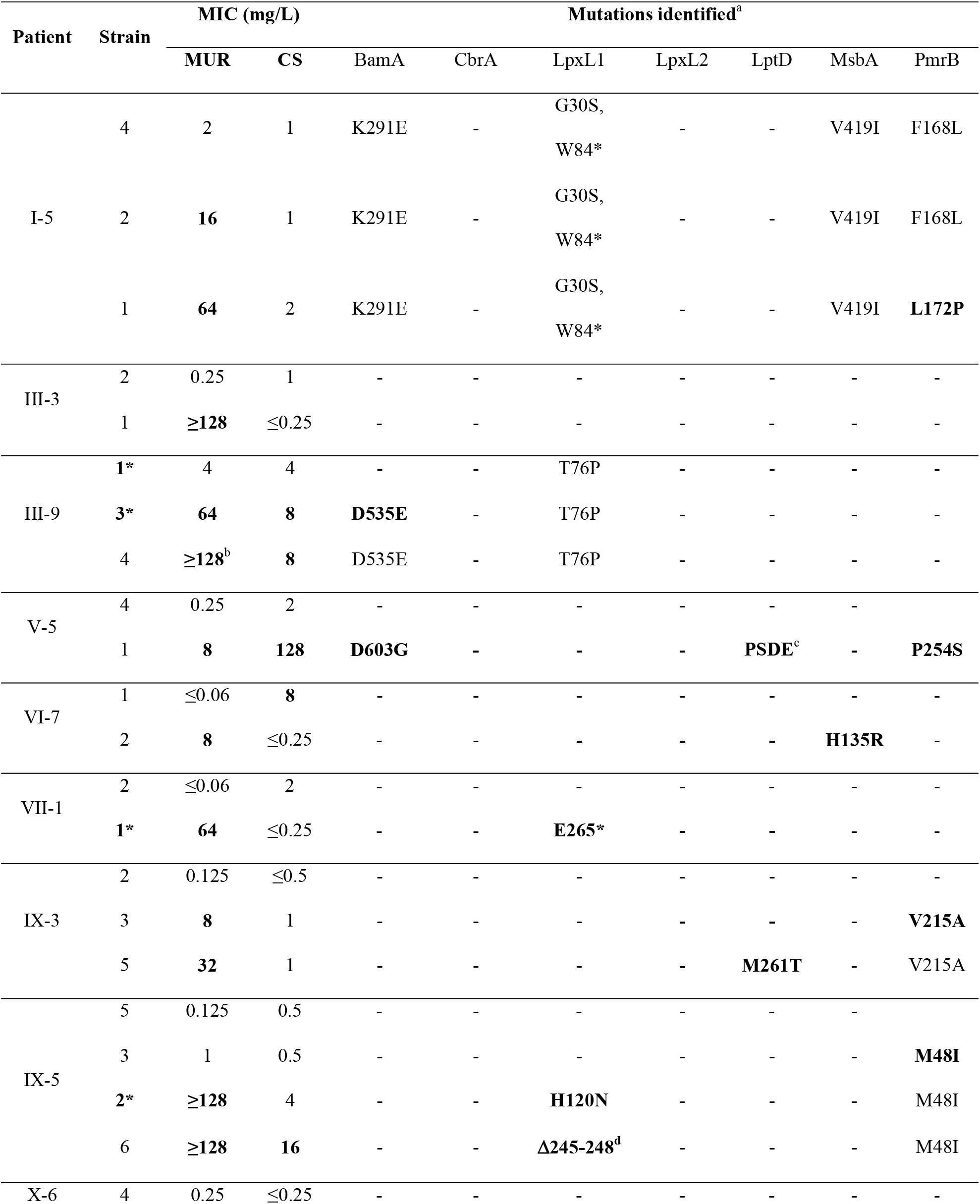
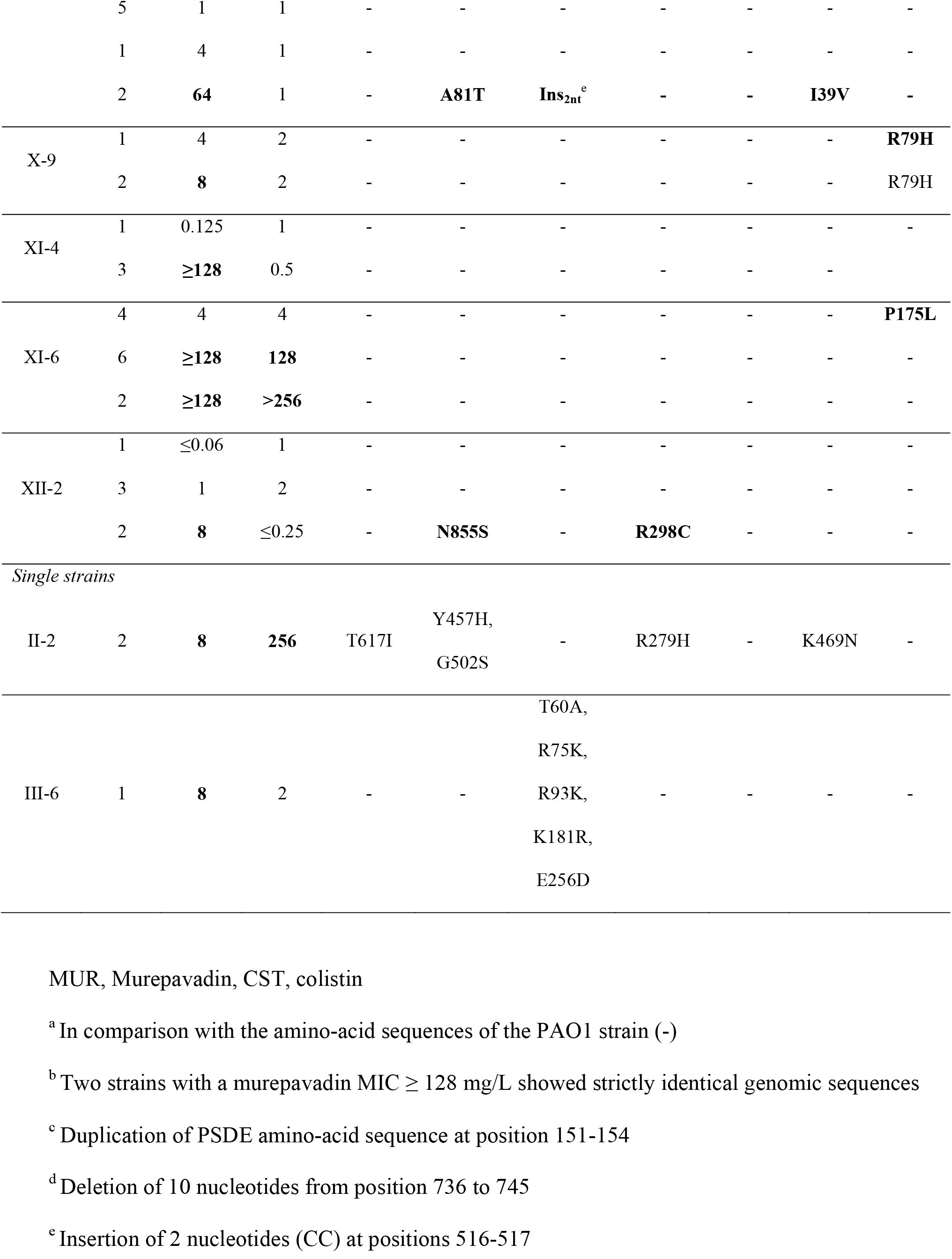
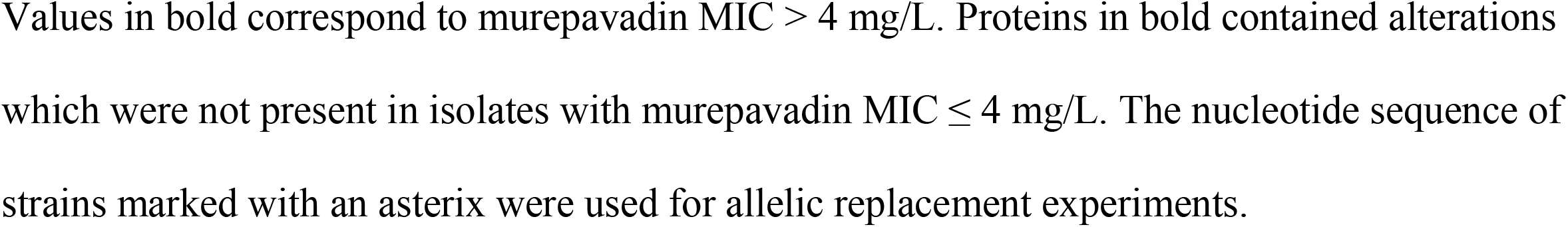
Protein or gene alterations identified at least two times in CF-strains with murepavadin MIC > 4 mg/l.

### Mutations in genes lpxL1 and bamA impact the activity of murepavadin in P. aeruginosa CF strains

Compared with their more susceptible counterparts, isolates with a murepavadin resistance > 4 mg/L (*i*.*e*., ≥ 128-fold the modal MIC for the whole population) displayed diverse SNPs in genes *bamA* (*n* =3 strains), *cbrA* (*n* = 3), *lpxL1* (*n* = 4), *lpxL2* (*n* = 1), *lptD* (*n* = 2), *msbA* (*n* = 2) and/or *pmrB* (*n* = 4) (Table 2). Genes *cbrA* and *pmrB* encode the sensor histidine kinases of two-component systems CbrA-CbrB, and PmrA-PmrB, respectively while the other loci are involved in the transport (*msbA, lptD, bamA*) or biosynthesis of LPS (*lpxL1, lpxL2*) (21). A first analysis of the distribution of these mutations among the selected strains failed to establish a correlation between murepavadin MICs and the alteration of specific genes or the number of mutated genes per isolate, suggesting that in CF strains murepavadin resistance is multifactorial and likely involves still unidentified loci. Supporting this assumption, several strains turned out not to harbor alterations in the short list of genes cited above, such as III-3-1, XI-4-3, XI-6-2, and XI-6-6 (murepavadin MIC ≥ 128mg/L, Table 2).

Genes *lpxL1* (synonym of *htrB1*) and *lpxL2* (*htrB2*) encode lauryl transferases known to modify the structure of lipid A in a site-specific manner. While LpxL1 mediates the addition of 2-hydroxylaurate at the C-2 position of lipid A, LpxL2 adds laurate at C-2’(22). Defects in either gene result in an increased permeability of the outer membrane, and hypersusceptibility to various antibiotics and polycationic peptides (23). Various patient-specific mutations were noted in the LpxL1-encoding gene resulting in either truncated peptides (W84*, E265*), amino acid substitutions (G30S, T76P, H120N, T60A, R75K, R93K, K181R, E256D), or disruption of the gene *lpxl1* itself (ins_2nt_ 516-517). To assess the impact of some of these alterations on murepavadin susceptibility levels, we replaced the *lpxL1* gene of strain PAO1 with the mutated alleles from clinical strains III-9-1 (inferred amino acid variation T76P), VII-1-1 (E265*) and IX-5-2 (H120N), respectively. These changes resulted in an 8-fold increase in murepavadin resistance (from 0.06 to 0.5 mg/L) (Table 3), in agreement with the reported emergence of a resistant *lpxL1* disruption mutant (MIC > 16 mg/L) along with several *lpxL2* mutants from hypermutator strain PAO1Δ*mutS* during time-kill experiments (15). Though it has been suggested that production of penta-acylated LPS molecules in LpxL1-deficient mutants results in decreased susceptibility to polymyxins, our results did not confirm such effects (Table 3) (23). In addition to *lpxL1*, the *bamA* gene was found to contain various single point mutations in a subset of CF strains exhibiting diverse resistance levels to murepavadin (from 2 to ≥ 128 mg/L), including the CF strain III-9-3 (Table 2). Its product, the β-barrel outer membrane protein BamA, is a component of the BAM complex (BamA-E). This complex plays an essential role in the folding of β-barrel proteins such as LptD and their insertion into the outer membrane (9). All the mutations identified in the selected CF strains (K291E, D535E, D603G, and T617I) were mapped in the β-strands of the C-terminal β-barrel domain. To assess the impact of the D535E substitution on murepavadin susceptibility, we first replaced the wild-type *bamA* gene of PAO1 with the mutated allele of strain III-9-3. This only resulted in a modest 2-fold increase in the peptide MIC (0.125 mg/L). However, the double replacement of *lpxL1* and *bamA* genes in the reference strain with the mutated alleles from strains III-9-1 and -3 that code for the T76P and D535E variations, respectively, had multiplicative effects on resistance to murepavadin (*e*.*g*., 32-fold increase as compared with the wild-type parent) (Table 3). The fact that the resistance of this double mutant (2 mg/L) is far below that of some CF strains (Table 2) reinforces the notion that other loci contribute to higher MIC values. Like with *lpxL1*, colistin susceptibility was unchanged in *bamA* mutants as compared with PAO1.

**TABLE 3.** Impact of the replacement of genes *lpxL1, bamA* and *pmrB* with mutated alleles on susceptibility of strain PAO1 to inhaled antibiotics.

| Strains | Alterations in proteins |  |  | MIC (mg/L) |  |  |
| --- | --- | --- | --- | --- | --- | --- |
|  | Lpxl1 | BamA | PmrB | Murepavadin | Colistin | Tobramycin |
| PAO1 wild-type | – | – | – | ≤ 0.06 | 0.5 | 0.5 |
| PAO1::*lpxL1*<sub>III-9-1</sub> | T76P | – | – | **0.5** | 0.5 | 0.25 |
| PAO1::*lpxL1*<sub>VII-1-1</sub> | E265* | – | – | **0.5** | 0.5 | 0.25 |
| PAO1::*lpxL1*<sub>IX-5-2</sub> | H120N | – | – | **0.5** | 0.5 | 0.25 |
| PAO1::*bamA*<sub>III-9-3</sub> | – | D535E | – | 0.125 | 0.5 | 0.5 |
| PAO1::*lpxL1*<sub>III-9-1</sub>-*bamA*<sub>III-9-3</sub> | T76P | D535E | – | **2** | 0.5 | 0.5 |
| PAO1Δ*pmrAB* | – | – | – | 0.03 | 0.5 | 0.25 |
| PAO1Δ*pmrAB*(pME6012) | – | – | – | 0.03 | 0.5 | 0.25 |
| PAO1Δ*pmrAB*(pAB3795) | – | – | G188D | **2** | **128** | **2** |
| PAO1Δ*prmAB*(pAB2243) | – | – | Q105P | **2** | **128** | **4** |
| PAO1Δ*pmrAB*(pAB3890) | – | – | D45E | **0.5** | **16** | **2** |

Values in bold correspond to MICs increased at least 4-fold as compared with those for wild-type PAO1

### Cross-resistance between tobramycin, colistin and murepavadin in pmrB clinical mutants

In addition to the genes involved in the transport and synthesis of LPS, single point mutations were identified in *pmrB* and *cbrA* leading to amino acid substitutions in their respective products, PmrB (M48I, R79H, F168L, L172P, P175L, V215A, P254S) and CbrA (A81T, Y457H, G502S, N855S). These histidine kinases sense and transmit stress signals from the cell envelope to their cognate cytoplasmic response regulators PmrA and CbrA, respectively, allowing for an appropriate adaptation of *P. aeruginosa*. Previous studies demonstrated that specific mutations in these phosphor-relays confer a dual resistance to polymyxins and aminoglycosides (24, 25). While inactivation of CbrA caused a modest augmentation of tobramycin (2-fold) and colistin MICs (4-fold), mutational activation of PmrB had much greater effects on the resistance to these cationic antibiotics (16-fold and 32-fold, respectively) (24, 25).

In the present strain collection, only one isolate (V-5-1) classified as colistin resistant (MIC = 128 mg/L) by reference to the EUCAST 2023 breakpoints, showed a mutation in sensor PmrB (P254S) (Table 2). To investigate on a possible PmrB-mediated cross-resistance to murepavadin, colistin and tobramycin, we selected eleven fully sequenced colistin-resistant *pmrB* mutants from the collection of the French National Reference Center for antibiotic resistance. As indicated in Table 4, these non-CF strains quite highly resistant to colistin (from 16 to > 256 mg/L) showed a susceptibility to murepavadin ranging from 0.25 to 8 mg/L. Reminiscent of the CF strains described in this study, multiple mutations in genes *bamA, cbrA, lptD, lpxL1* and *lpxL2*, were also present in these bacteria. Constructs of plasmid vector pME6012 carrying the *pmrAB* operons from three strains (3795, 2243, 3890) were used to complement the deletion mutant PAO1Δ*pmrAB*. Confirming the impact of PmrB amino-acid substitutions G188D, Q105P and D45E on the susceptibility to murepavadin, MIC values of the peptide increased from 16 to 64-fold upon complementation, thus reaching 0.5 and 2 mg/L, respectively (Table 3). Colistin MICs varied in parallel (from 32-fold to 256-fold) suggesting that the degree of resistance to both antibiotics is modulated by specific amino acid variations in different regions of PmrB (26). Of note, the G188D and Q105P substitutions are located in the HAMP and periplasmic domains of PmrB, respectively while D45E affects the periplasmic domain of the sensor. Consistent with our results, spontaneous *pmrB* mutants 8- to 32-fold more resistant than parental strain PA14 to murepavadin analogue POL7001 have been previously described (16). As shown by our laboratory, alteration of sensor PmrB can potentially be responsible for a decreased susceptibility of *P. aeruginosa* to aminoglycosides (up to 16-fold) (Table 3) (25). Since mutations may target the *pmrB* gene in the context of CF lung chronic colonization, the probability that some of them affect the activity of the three molecules, murepavadin, colistin and tobramycin, should be considered. Longitudinal studies enrolling cohorts of CF patients would be necessary to validate this hypothesis, looking at the emergence of cross-resistant mutants under murepavadin, colistin or tobramycin aerosol therapy.

**TABLE 4.**
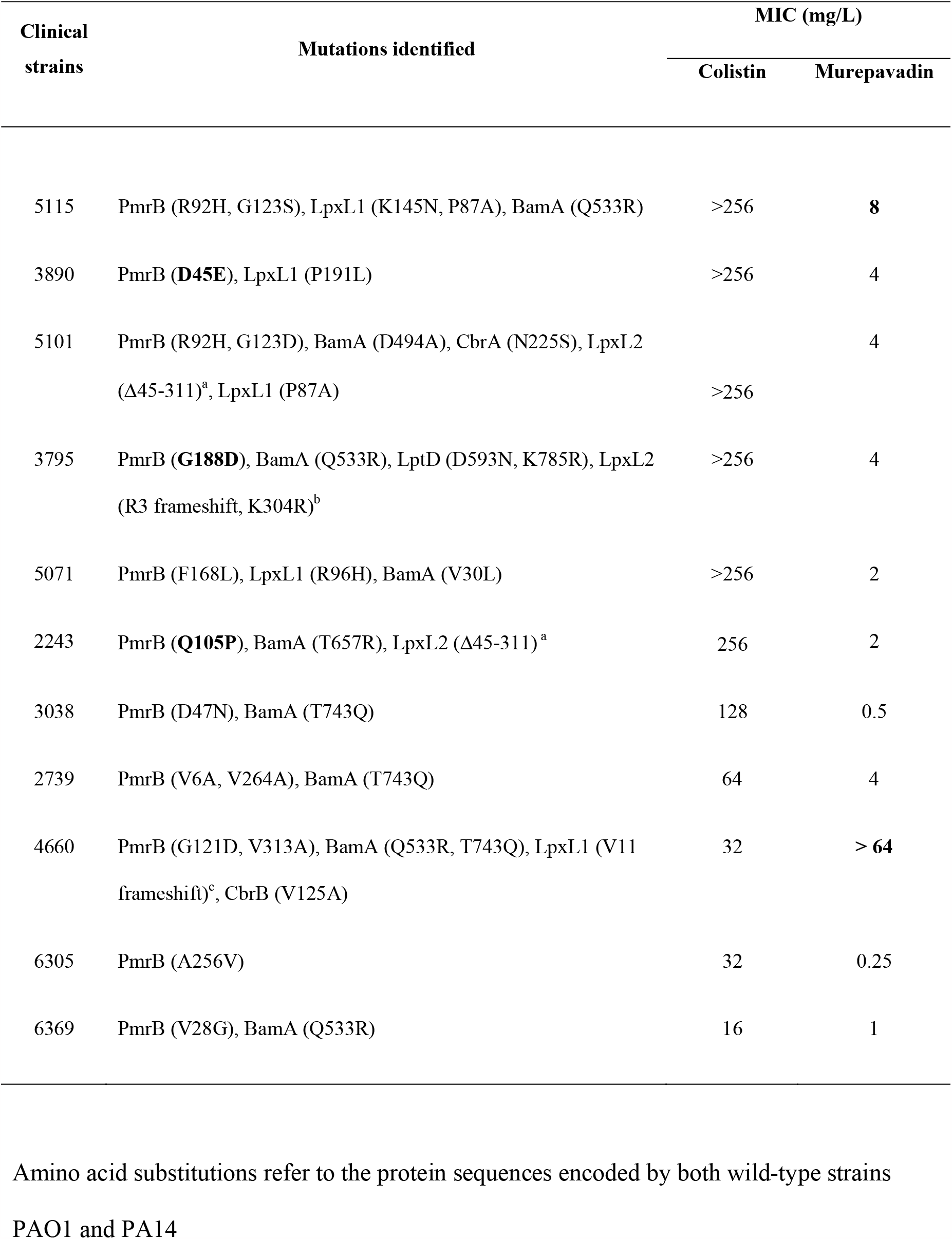
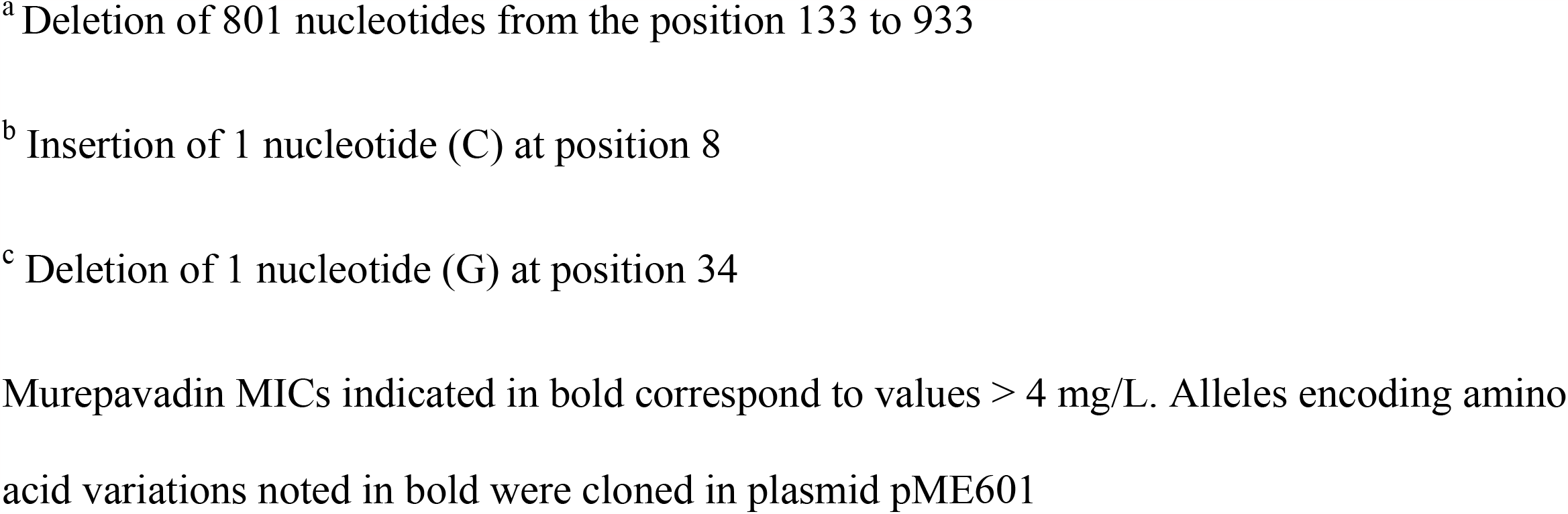
Murepavadin susceptibility of colistin-resistant clinical strains harboring mutations in gene *pmrB*.

| Clinical strains | Mutations identified | MIC (mg/L) |  |
| --- | --- | --- | --- |
|  |  | Colistin | Murepavadin |
| 5115 | PmrB (R92H, G123S), LpxL1 (K145N, P87A), BamA (Q533R) | >256 | **8** |
| 3890 | PmrB (**D45E**), LpxL1 (P191L) | >256 | 4 |
| 5101 | PmrB (R92H, G123D), BamA (D494A), CbrA (N225S), LpxL2 (Δ45-311)<sup>a</sup>, LpxL1 (P87A) | >256 | 4 |
| 3795 | PmrB (**G188D**), BamA (Q533R), LptD (D593N, K785R), LpxL2 (R3 frameshift, K304R)<sup>b</sup> | >256 | 4 |
| 5071 | PmrB (F168L), LpxL1 (R96H), BamA (V30L) | >256 | 2 |
| 2243 | PmrB (**Q105P**), BamA (T657R), LpxL2 (Δ45-311)<sup>a</sup> | 256 | 2 |
| 3038 | PmrB (D47N), BamA (T743Q) | 128 | 0.5 |
| 2739 | PmrB (V6A, V264A), BamA (T743Q) | 64 | 4 |
| 4660 | PmrB (G121D, V313A), BamA (Q533R, T743Q), LpxL1 (V11 frameshift)<sup>c</sup>, CbrB (V125A) | 32 | **> 64** |
| 6305 | PmrB (A256V) | 32 | 0.25 |
| 6369 | PmrB (V28G), BamA (Q533R) | 16 | 1 |

Amino acid substitutions refer to the protein sequences encoded by both wild-type strains
PAO1 and PA14
<sup>a</sup> Deletion of 801 nucleotides from the position 133 to 933
<sup>b</sup> Insertion of 1 nucleotide (C) at position 8
<sup>c</sup> Deletion of 1 nucleotide (G) at position 34
Murepavadin MICs indicated in bold correspond to values > 4 mg/L. Alleles encoding amino acid variations noted in bold were cloned in plasmid pME601

### Other mutations identified

Unexpectedly, mutations in the murepavadin target protein LptD were identified in only two CF strains that otherwise displayed multiple alterations in their DNA repair systems (4) (Tables 2 and Table S2). A tandem duplication of the PSDE sequence spanning from positions 151 to 154 was found in V-5-1, a strain with a murepavadin resistance equal to 8 mg/L that also harbored mutations in PmrB (P254S) and BamA (D603G). Though the impact of this structural change on the function of LptD was not investigated further, it is interesting to note that a tandem duplication of residues LRDKGM at positions 210 to 215 together with a G214D change was reported previously for an *in vitro* selected mutant showing a 64-fold higher murepavadin resistance than its parent PAO1 (4, 15). The second isolate (IX-3-5) of this study displaying a LptD variant (M261T) contained a concomitant V215A change in PmrB, for a resistance level to murepavadin equal to 32 mg/L. Again, highlighting the multifactorial nature and complexity of mechanisms contributing to elevated MICs of the peptide, 511 SNPs were identified between the susceptible isolate IX-3-3 (MIC = 0.125 mg/L) and its counterpart, IX-3-5 (Table S2).

ATPase MsbA is a member of the ABC-transporter superfamily. The role of this transmembrane protein is to flip complete lipid A-core molecules from the inner to the outer side of the cytoplasmic membrane before their modification and subsequent transport by the Lpt machinery to the cell surface (27). Five CF strains of our collection produced MsbA proteins with single amino acid substitutions (V419I in I-5-4, I-5-2, I-5-1; H135R in VI-7-2; I39V in X-6-2). In contrast to the other isolates, strain VI-7-2 did not appear to contain alterations in genes *bamA, cbrA, lptD, lpxL1, lpxL2* and *pmrB*. Interestingly, its genetic divergence from its susceptible counterpart VI-7-1 (MIC ≤ 0.06 mg/L) was limited to 26 SNPs, which suggests a contribution of the H135R mutation to the resistance of VI-7-2 to murepavadin (8 mg/L). Little is known about the impact the alteration of MsbA may have on the fitness of *P. aeruginosa*. In *Escherichia coli*, experiments demonstrated that the deletion of the MsbA-encoding gene drastically decreases cell viability both *in vitro* and *in vivo* (28). However, amino acid substitutions in the protein, which shares 40.3% sequence identity with its homologue in *P. aeruginosa*, can be tolerated as those conferring resistance to quinoline compounds targeting MsbA (28). Further experiments are required to clarify to which extent mutations in MsbA and LptD reduce the susceptibility of CF strains to murepavadin and may affect the fitness of *P. aeruginosa*.

## Conclusion

The present study confirms the good *in vitro* activity of murepavadin on CF strains, making this original antipseudomonal peptide endowed with a unique mode of action an interesting alternative to older antibiotics currently used by inhalation, such as colistin, tobramycin and aztreonam. However, although murepavadin is still under development, a notable proportion of the *P. aeruginosa* strains that already colonize CF patients display various degrees of resistance to the drug, including some isolates for which MICs are > 4 mg/L (9.1%) (*i*.*e*., ≥ 128-fold the modal value of our strain population). It is now well established that long-term and repeated administration of antibiotics to CF patients select bacterial subpopulations increasingly resistant to one or more antimicrobials (29). A trivial explanation for the presence of *P. aeruginosa* relatively resistant to murepavadin in patients never treated with this drug could be that current treatments by aminoglycosides (mostly tobramycin) and/or polymyxins (mostly colistin) select mutation-based mechanisms of cross-resistance implying global regulators or two-components such as PmrAB. Aminoglycosides, polymyxins and murepavadin have in common to interact with components of the bacterial outer membrane. Thus, it is tempting to speculate that still unknown mechanisms impairing the activities of these antibiotic families are related to the structure or physiology of the cell envelope, as for the resistant strains from patients III-3, X-6, XI-4 and XI-6. However, it remains unclear which selective pressure in the CF lung can lead to the emergence of mutants specifically resistant to murepavadin while the drug has never been used (*e*.*g, lpxL1*). Mutants exhibiting various modifications in the structure of the lipid A have already been reported in the context of CF, which could reflect a phenotypic adaption of *P. aeruginosa* to this particular lung environment, not necessarily linked to the presence of antibiotics (30). Understanding the phenotypic and genotypic evolution of CF strains under murepavadin therapy will be key to the positioning of this new agent among the antibiotic resources available against *P. aeruginosa*.

## Materials and methods

### Bacterial strains, culture media, and growth conditions

The strains and plasmids used in this study are described in Table S3. During a four-month multicenter national survey (GERPA MUCO II, from October 2019 to January 2020) involving twelve French hospitals (Besançon, Brest, Limoges, Lyon La Croix Rousse, Nantes, Paris Foch, Paris Necker, Paris Robert Debré, Toulon, Toulouse, Reims and Rennes), 718 isolates of *P. aeruginosa* were collected from 120 CF patients (ten patients per participating center, and six colonies randomly picked from a single sputum sample per individual). A subcollection of 230 strains from 105 CF patients aged 1 to 52-years (median 24-years) was selected for the present study, to retain only those isolates exhibiting different antibiotic susceptibility profiles (at least a 2-fold MIC difference for at least 2 antibiotics, data not shown). The collection was enriched with 11 non-CF colistin-resistant clinical strains of *P. aeruginosa* (MIC > 4 mg/L) harboring a PmrB mutation, isolated between 2014 and 2019 in eleven French hospitals (repository of the French National Reference Center for antibiotic resistance, Besançon hospital). All strains were grown at 35 +/-2°C in Mueller-Hinton broth (MHB) (Dickinson Microbiology Systems, Cockeysville, Md, United States) with adjusted concentrations of divalent cations Ca^2+^ and Mg^2+^ or on Mueller-Hinton Agar (MHA) plates (Bio-Rad, Marnes-la-Coquette, France). In conjugation experiments, transconjugants were selected on *Pseudomonas* Isolation Agar (PIA, Becton Dickinson) supplemented with 2,000 mg/L streptomycin. The plasmid has been excised after culture of the transconjuguants on a M9 minimal medium (42 mM Na_2_HPO_4_, 22 mM KH_2_PO_4_, 19 mM NH_4_Cl, 8.5 mM NaCl) with 5% sucrose as a source of carbon and energy.

### Antimicrobial susceptibility testing

Minimum inhibitory concentrations (MICs) of ticarcillin, piperacillin plus 4 mg/L tazobactam, aztreonam, ceftazidime, ceftolozane plus 4 mg/L tazobactam, ceftazidime plus 4 mg/L avibactam, cefepime, imipenem, meropenem, amikacin, tobramycin and ciprofloxacin were determined in MHB by using customized microplates containing lyophilized antibiotic powders (Thermo Fisher, Illkirch-87 Graffenstaden, France). MICs of colistin (from 0.12 to 256 mg/L) and murepavadin (from 0.06 to 128 mg/L) were determined by the standard microdilution method in MHB using titrated powders of colistin sulfate (Sigma-Aldrich) and murepavadin (ProbeChem, China) (31). The strains were categorized as susceptible (S), susceptible at increased exposure (I) or resistant (R) according to the EUCAST 2023 clinical breakpoints (32). Quality controls in MIC experiments were performed on a regular basis with *P. aeruginosa* strains ATCC 27853 and PAO1, and *E. coli* NCTC 13846.

### SNP identification

Twenty-one CF strains with a murepavadin MIC value > 4 mg/L were submitted to complete genome sequencing along with 17 more susceptible isolates (≤ 4 mg/L) coexisting in the same sputum samples. Whole DNA was extracted from overnight cultures by using the PureLink Genomic DNA mini kit (Thermo Fisher Scientific). Libraries were prepared (Nextera XT DNA Library Preparation kit) and sequenced on an Illumina NextSeq 500 platform (Illumina, San Diego, CA; P2M platform, Institut Pasteur, Paris, France). Fastq files were generated and demultiplexed with bcl2fastq Conversion Software (v2.20; Illumina). The final average sequencing depth was > 80 X for all of the strains. The reads were assembled using Shovill-Spades (v3.14.0) and the contigs annotated with Prokka (v1.14.5). Single Nucleotide Polymorphisms (SNPs) (accession number NC_002516.1) were detected by mapping the reads against the reference strain PAO1 sequence, by using BioNumerics (v7.6.3) software (Applied Maths) with a minimum sequencing depth of 10 X. Sequence Types (STs) were determined according to the MLST scheme available at PubMLST (https://pubmlst.org).

### Allelic replacement

Respective sequences of genes *lpxL1* (888-bp in length) and *bamA* (2,394-bp) were amplified by PCR from whole DNA extracts of strains III-9-1, III-9-3, VII-1-1 and IX-5-2 (Genomic DNA extraction kit, Macherey-Nagel, Hoerdt) by using the primers listed in Table S4. The resultant fragments were cloned into plasmid pKNG101 by using the NEBuilder® HiFi DNA Assembly Cloning kit (New England Biolabs, Ipswich, MA, USA) (33). The recombinant plasmids were next transferred to *E. coli* CC118λ*pir* by transformation and then to strain PAO1 by triparental mating with helper strain *E. coli* HB101(pRK2013) (34). Transconjugants were selected on PIA medium containing 2,000 mg/L streptomycin. Excision of integrated plasmids was obtained by replica plating on M9 minimal agar medium supplemented with 5% sucrose. The allelic replacement of genes *lpxL1* and *bamA* in PAO1 was checked PCR sequencing (RUO3500 Genetic Analyzer, Applied Biosystems) with specific primers (Table S4).

## Acknowledgements

We thank Emma Girardin and Jade Chaillon for their excellent technical assistance. We are grateful to the members of the MucoMicrobes study group for collecting strains: Emilie Cardot-Martin (Centre Hospitalier Universitaire Foch, Paris), Vincent Cattoir (Centre Hospitalier Universitaire de Rennes), Lise Crémet (Centre Hospitalier Universitaire de Nantes), Anne Doléan-Jordheim (Hospices civils de Lyon), Agnès Ferroni (Centre Hospitalier Universitaire de Necker, Paris), Fabien Garnier (Centre Hospitalier Universitaire de Limoges), Hélène Guet-Revillet (Centre Hospitalier Universitaire de Toulouse), Thomas Guillard (Centre Hospitalier Universitaire de Reims), Geneviève Hery-Arnaud (Centre Hospitalier Universitaire de Brest), Guenièvre Imbert (Centre Hospitalier de Toulon), and Patricia Mariani (Centre Hospitalier Universitaire Robert Debré, Paris).

## Transparency declarations

None to declare.

